# An inhibitor/anti-inhibitor system controls the activity of lytic transglycosylase MltF in *Pseudomonas aeruginosa*

**DOI:** 10.1101/2023.07.28.551027

**Authors:** Michelle Wang, Sheya Xiao Ma, Andrew J. Darwin

**Affiliations:** Department of Microbiology, New York University Grossman School of Medicine, New York, New York, United States

## Abstract

Most bacterial cell envelopes contain a cell wall layer made of peptidoglycan. The synthesis of new peptidoglycan is critical for cell growth, division and morphogenesis, and is also coordinated with peptidoglycan hydrolysis to accommodate the new material. However, the enzymes that cleave peptidoglycan must be carefully controlled to avoid autolysis. In recent years, some control mechanisms have begun to emerge, although there are many more questions than answers for how most cell wall hydrolases are regulated. Here, we report a novel cell wall hydrolase control mechanism in *Pseudomonas aeruginosa*, which we discovered during our characterization of a mutant sensitive to the overproduction of a secretin protein. The mutation affected an uncharacterized Sel1-like repeat protein encoded by the PA3978 locus. In addition to the secretin-sensitivity phenotype, PA3978 disruption also increased resistance to a β-lactam antibiotic used in the clinic. *In vivo* and *in vitro* analysis revealed that PA3978 binds to the catalytic domain of the lytic transglycosylase MltF and inhibits its activity. ΔPA3978 mutant phenotypes were suppressed by deleting *mltF*, consistent with them having been caused by elevated MltF activity. We also discovered another interaction partner of PA3978 encoded by the PA5502 locus. The phenotypes of a ΔPA5502 mutant suggested that PA5502 interferes with the inhibitory function of PA3978 towards MltF, and we confirmed that activity for PA5502 *in vitro*. Therefore, PA3978 and PA5502 form an inhibitor/anti-inhibitor system that controls MltF activity. We propose to name these proteins Ilt (inhibitor of lytic transglycosylase) and Lii (lytic transglycosylase inhibitor, inhibitor).

**IMPORTANCE:** A peptidoglycan cell wall is an essential component of almost all bacterial cell envelopes, which determines cell shape and prevents osmotic rupture. Antibiotics that interfere with peptidoglycan synthesis have been one of the most important treatments for bacterial infections. Peptidoglycan must also be hydrolyzed to incorporate new material for cell growth and division, and to help accommodate important envelope-spanning systems. However, the enzymes that hydrolyze peptidoglycan must be carefully controlled to prevent autolysis. Exactly how this control is achieved is poorly understood in most cases, but is a highly active area of current research. Identifying hydrolase control mechanisms has the potential to provide new targets for therapeutic intervention. The work here reports the important discovery of a novel inhibitor/anti-nhibitor system that controls the activity of a cell wall hydrolase in the human pathogen *Pseudomonas aeruginosa*, and which also affects resistance to an antibiotic used in the clinic.

## INTRODUCTION

The Gram-negative bacterium *Pseudomonas aeruginosa* occurs widely in the environment and can cause serious opportunistic human infections (1). *P. aeruginosa* is one of a small group of pathogens causing the majority of hospital-acquired infections, and The World Health Organization has declared it a “Priority 1: Critical” pathogen (2, 3). Increasing antibiotic resistance, including outbreaks of multidrug resistant strains in hospitals, emphasizes the need for continued study and the development of effective treatments (1, 4). The bacterial cell envelope is critical for interaction with the host, for export/assembly of virulence factors, and it is a target for antimicrobial agents. As for almost all bacteria, the *P. aeruginosa* cell envelope contains a mesh-like cell wall layer composed of peptidoglycan. Peptidoglycan consists of linear glycan strands of alternating *N*-acetyl muramic acid and *N*-acetyl glucosamine disaccharide units, with a short amino acid peptide chain attached to *N*-acetyl muramic acid (5, 6). Covalent bonds between peptide chains of adjacent glycan strands result in an extensively cross-linked cell wall that withstands turgor pressure. New peptidoglycan must be synthesized for cell elongation and division, but at the same time, existing peptidoglycan must be hydrolyzed to accommodate the incorporation of new material (7, 8). Peptidoglycan hydrolysis is also needed for bacterial morphogenesis and for the insertion of trans-envelope structures such as secretion systems and flagella (9–11).

Most bacteria encode numerous hydrolases that can cleave all or almost all of the different bonds within intact peptidoglycan, and in some of the resulting soluble products (9–11). However, cleaving peptidoglycan comes with the potential for catastrophic consequences if it is not carefully controlled. Uncovering the mechanisms by which these hydrolase enzymes are controlled is an especially active area of research, which has the potential to identify new therapeutic targets in pathogenic species. For many peptidoglycan hydrolases, there is either no information, or incomplete information about how their activity is regulated, although some control mechanisms have begun to emerge in recent years (see ref. 12 for a recent review). These mechanisms include modification of the peptidoglycan substrate, direct activation or inhibition by trans-acting proteins or other molecules, and the control of hydrolase concentration by various mechanisms. One mechanism by which some peptidoglycan hydrolases are controlled is through their degradation by a carboxyl-terminal processing protease (CTP; 13, 14).

Our laboratory came across the *P. aeruginosa* CTP CtpA during a screen for sensitivity to the overproduction and mislocalization of a pore-forming outer membrane secretin protein (15, 16). Secretins are critical components of type 2 and 3 secretion systems, and type IV pili (17–19). However, they can mislocalize into the inner membrane, which is lethal unless the phage shock protein (Psp) stress response mitigates it in some species (reviewed in ref. 20). *P. aeruginosa* produces multiple secretins, but it does not have a Psp response system (21). This provided our original motivation to identify mutations that caused secretin-sensitivity, one of which disrupted *ctpA*. CtpA is also required for normal function of the type 3 secretion system and for virulence in a mouse model of acute pneumonia (15). Our characterization of CtpA revealed that it controls the activity of peptidoglycan cross-link hydrolases by degrading at least four of them (14). We do not yet know the molecular mechanism that explains the secretin-sensitivity phenotype of a *ctpA* null mutant, but the finding that CtpA controls peptidoglycan hydrolases might provide a hint. Secretins must navigate through the cell wall on their way to the outer membrane, and so altered peptidoglycan architecture might increase toxicity by disturbing secretin trafficking. Or perhaps altered peptidoglycan metabolism affects phospholipid and lipopolysaccharide biosynthesis, which share precursors in common with peptidoglycan, rendering the cell envelope more susceptible to the stress.

Our finding that CtpA is involved in cell wall hydrolase control has motivated us to reexamine other genes we found in our secretin sensitivity screen. We were especially interested in genes encoding cell envelope proteins of unknown function. One of these is PA3978, a predicted periplasmic Sel1-like repeat protein that has not been characterized. In this study, we report that our investigation of this protein has revealed a novel example of cell wall hydrolase control. PA3978 is a direct inhibitor of the lytic transglycosylase MltF, an enzyme that cleaves the glycan chains of peptidoglycan. We also discovered that PA3978 is acted on by another previously uncharacterized envelope protein, which acts as an anti-inhibitor of PA3978.

## RESULTS

### Identification of a PA3978 null mutant

Outer membrane pore-forming secretin proteins can mislocalize and cause cell envelope toxicity. To investigate how this toxicity might be counteracted in *P. aeruginosa*, we previously reported the isolation of transposon insertion mutants sensitive to increased production of the XcpQ secretin (16). One insertion was in *ctpA*, which encodes a carboxyl-terminal processing protease that degrades cell wall hydrolases (14, 15, 22–24). Recently, we have investigated some genes of unknown function that were discovered in the same screen that found *ctpA*. One mutant had a transposon insertion in the PA3978 gene (PAO1 strain designation), which encodes a predicted soluble 17 kDa periplasmic protein after removal of a type I signal sequence. The arrangement of the PA3978 gene makes polar effects unlikely (Fig. 1a). However, to confirm that the XcpQ-sensitivity was not caused by an unlinked spurious mutation, the transposon insertion was backcrossed into the wild type strain, which reproduced the XcpQ-sensitive phenotype (Fig. 1b). We also constructed a ΔPA3978 in frame deletion mutant, which had the same XcpQ-sensitivity as the transposon insertion mutant (Fig.1b). Finally, plasmid-encoded PA3978 was toxic, especially in the wild type strain that also had the endogenous protein. Nevertheless, this plasmid was able to complement the reduced growth phenotype of a ΔPA3978 mutant during XcpQ overproduction (Supplemental Fig. S1).

**FIG 1.**
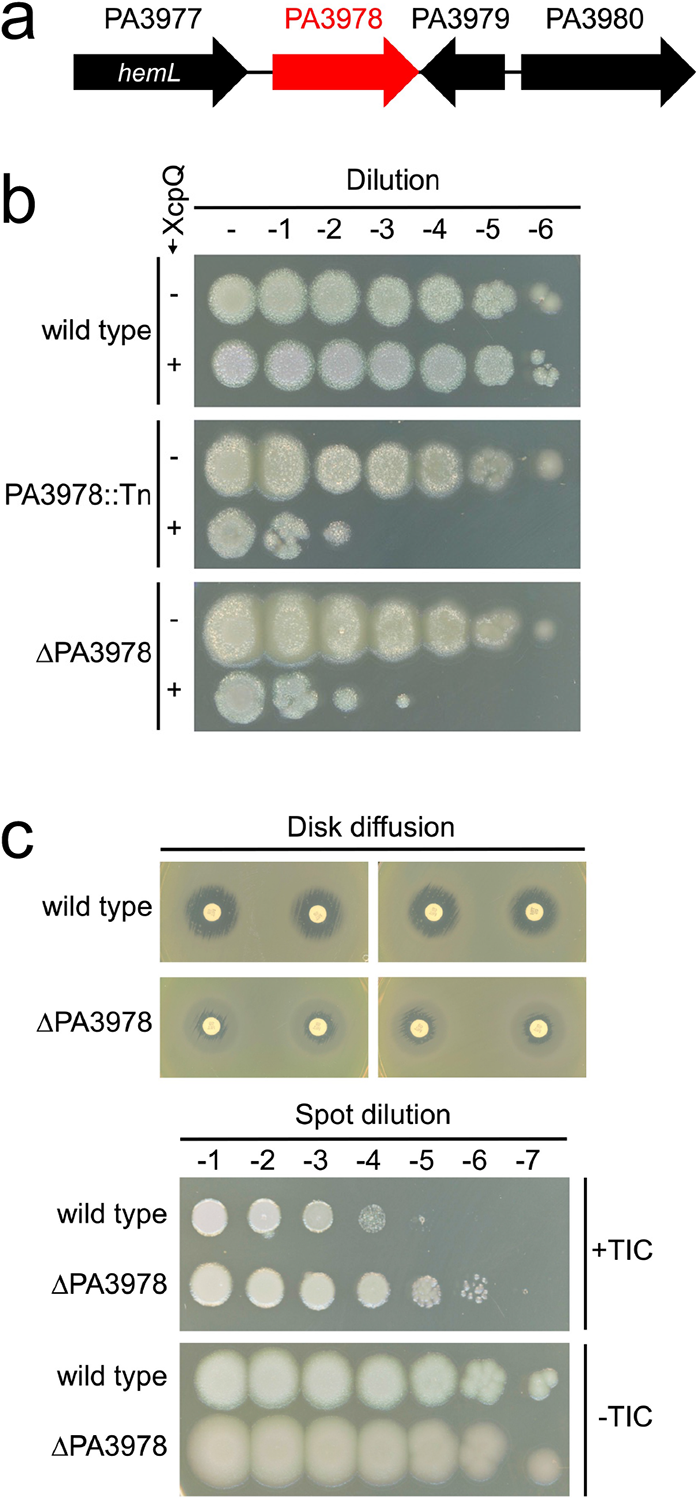
PA3978 null mutant phenotypes. (a) Organization of the PA3978 locus. Gene numbers are from strain PAO1. Only PA3977 has a previously assigned gene name. (b) Sensitivity to XcpQ secretin overproduction. Each strain contained either the *tac* promoter expression plasmid pVLT35 (-XcpQ), or the *xcpQ*^+^ derivative pAJD942 (+XcpQ). Serial dilutions of normalized saturated cultures were spotted onto LB agar containing IPTG and incubated at 37°C for approximately 24 h. (c) Top, disc diffusion assays: discs containing Ticarcillin-clavulanic acid (75/10 µg) were placed onto Mueller-Hinton agar that had been inoculated with a lawn of the indicated strain (duplicate experiments are shown, with two discs used in each one). Bottom, plating efficiency assay: serial dilutions of normalized saturated cultures were spotted onto the surface of Mueller-Hinton agar with (+TIC) or without (-TIC) 4 µg/ml ticarcillin-clavulanate (15:1). For both assays, plates were incubated at 37°C for approximately 24 h. PA3978::Tn is the original transposon insertion mutation backcrossed into the wild type and ΔPA3978 is an in frame deletion mutant.

An imipenem-sensitive PA3978 transposon insertion mutant was isolated by others in a screen for strain PA14 mutants with altered susceptibility to β-lactam antibiotics (25). Our PAK ΔPA3978 mutant did not have obvious imipenem sensitivity in a disk diffusion assay screen (data not shown). However, we did discover a resistance phenotype for the clinically relevant β-lactam/β-lactamase inhibitor combination ticarcillin-clavulanate (TIC; Fig. 1c). We developed a sensitive and highly reproducible spot-dilution plating efficiency assay on agar containing 4 µg/ml TIC, which proved valuable to monitor this phenotype with high resolution in subsequent genetic experiments (Fig. 1c and below). β-lactam antibiotics inhibit peptidoglycan synthesis, which suggested the possibility that PA3978 might have a link to peptidoglycan metabolism. Our subsequent experiments confirmed this possibility.

### PA3978 interacts with MltF and PA5502

The AlphaFold-predicted structure of PA3978 is almost entirely α-helical (Fig. 2a). Its primary sequence has three Sel1-like repeat domains predicted by TPRpred, located at amino acids 26-61, 62-97 and 98-133, with P-values of 1.7e-03, zero and 1.3e-12, respectively (Fig. 2a; ref. 26). Sel1-like repeats are a subtype of tetratricopeptide repeats (TPR) that mediate protein-protein interactions (27, 28). Therefore, it seemed likely that PA3978 interacted with one or more proteins. To test this hypothesis, three independent purifications of PA3978 with a C-terminal FLAG tag were analyzed by mass spectrometry (MS), along with three negative control purifications in which the PA3978 bait protein was not FLAG-tagged. There were 67 known or predicted envelope proteins present in all three PA3978-FLAG purifications (Supplemental Table S1). Comparison with the levels in the negative controls revealed that two proteins were both abundant and highly enriched with PA3978-FLAG: the lytic transglycosylase MltF (PA3764) and a predicted outer membrane lipoprotein of unknown function, PA5502 (Fig. 2b).

**FIG 2.**
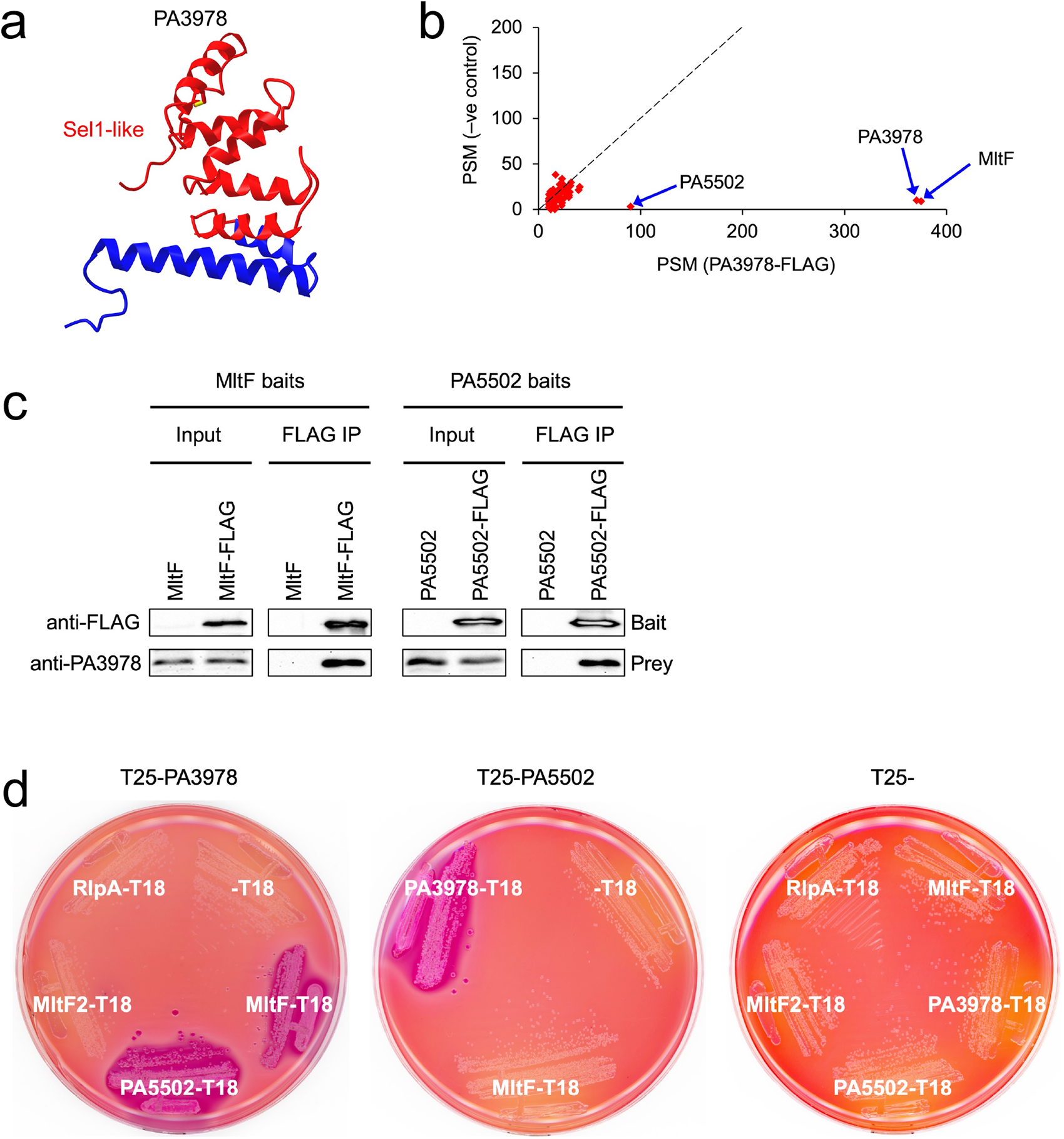
Discovery and verification of PA3978 binding partners. (a) AlphaFold-predicted structure of the mature PA3978 protein from the AlphaFold Protein Structure Database. The region encompassing the three Sel1-like repeats is colored in red and the remaining C-terminal region in blue. (b) Scatter plot of proteins identified by mass spectrometry after purification from strains encoding C-terminal FLAG-tagged PA3978 or untagged PA3978. Data are the average peptide spectrum matches (PSM) from triplicate experiments. The dotted line represents the theoretical position of proteins of equal abundance in both samples based on PSM. (c) Reciprocal pull down verification assay. Immunoblot analysis of input lysates and immunoprecipitates (FLAG IP). Baits were MltF-FLAG or MltF negative control (left) and PA5502-FLAG or PA5502 negative control (right). Immunoblots are single representatives of several independent replicate experiments. (d) Bacterial two-hybrid analysis. *E. coli* BTH101 strains grown on MacConkey-maltose agar contained a plasmid encoding PA3978 fused to the C-terminus of Cya-T25 (left), PA5502 fused to the C-terminus of Cya-T25 (middle), or Cya-T25 only (right). Strains contained a second plasmid encoding different proteins fused to the N-terminus of Cya-T18 as indicated.

To verify the mass spectrometry findings for MltF and PA5502, we purified MltF-FLAG and PA5502-FLAG bait proteins and confirmed that endogenous PA3978 co-purified with each by immunoblot (Fig. 2c). We also used a bacterial two-hybrid assay (BACTH), which tests for proximity of two fragments (T18 and T25) of the catalytic domain of *Bordetella pertussis* adenylate cyclase (29). Proteins fused to T18 and T25 restore Cya activity if they associate. In an *E. coli* Δ*cya* mutant this activates cAMP-CRP-dependent genes such as those required for maltose catabolism, producing red colonies on MacConkey-maltose agar. This analysis supported robust and independent PA3978-MltF and PA3978-PA5502 interactions (Fig. 2d). BACTH analysis also suggested that MltF and PA5502 do not interact with each other (Fig. 2d). We tested the latter conclusion further by analyzing all possible pairwise combinations of MltF and PA5502 N- and C-terminal T18 or T25 fusion proteins, and all were negative (data not shown). In agreement with the MS analysis, BACTH experiments also suggested that PA3978 is not a general interactor with lytic transglycosylases, as it tested negative for interaction with lytic transglycosylases RlpA (PA4000) and also MltF2 (PA2865), a homolog of MltF (Fig. 2d).

### Deletion of *mltF* suppresses ΔPA3978 mutant phenotypes

We first focused on the implications of the PA3978-MltF interaction, because MltF is a lytic transglycosylase, and MltF co-purified with similar abundance to PA3978-FLAG based on peptide spectral matches (Fig. 2b; Supplemental Table S1; 30, 31). We tested for a functional relationship between PA3978 and MltF by comparing the phenotypes of strains with ΔPA3978 and/or Δ*mltF* in frame deletion mutations (Fig. 3).

**FIG 3.**
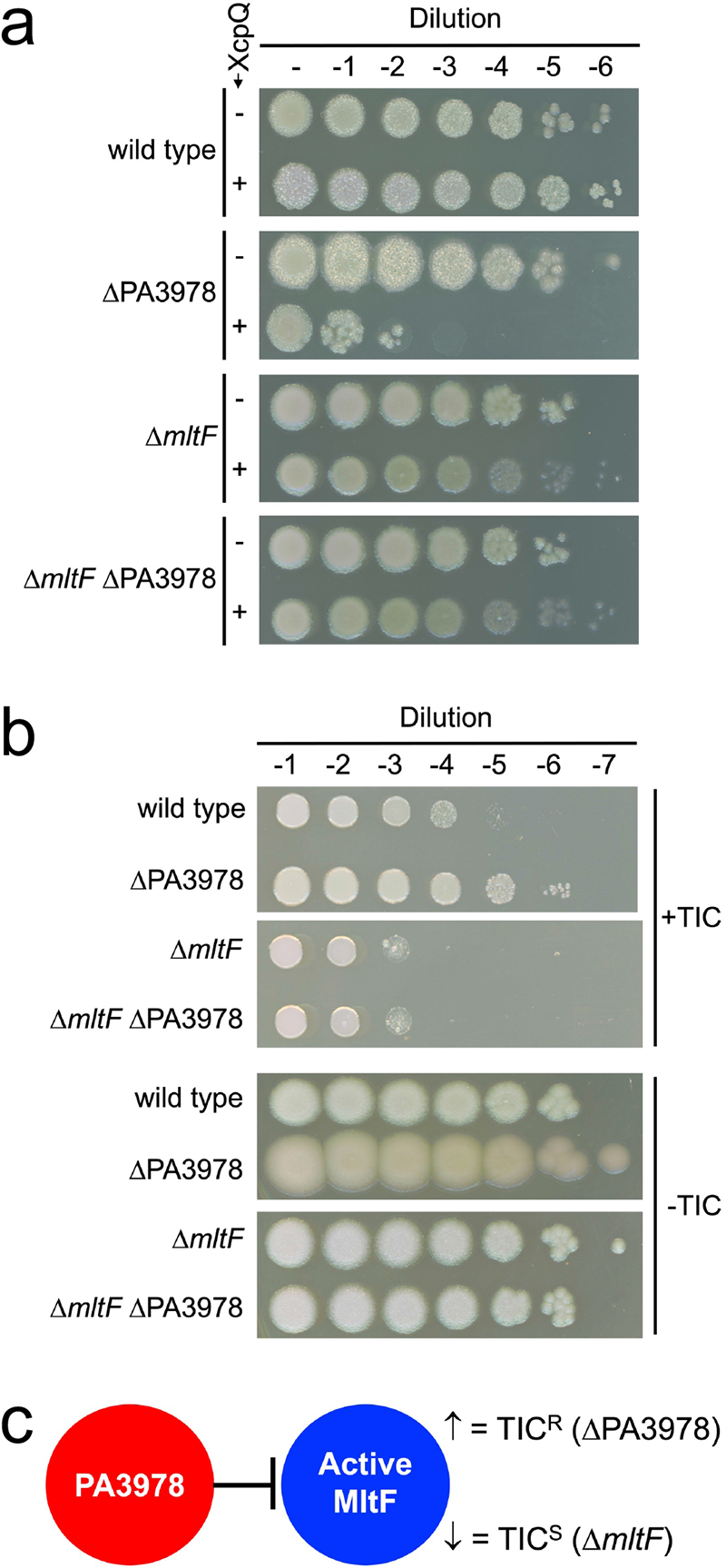
Δ*mltF* mutant phenotypes. (a) Sensitivity to XcpQ secretin overproduction. Each strain contained either the *tac* promoter expression plasmid pVLT35 (-XcpQ), or the *xcpQ*^+^ derivative pAJD942 (+XcpQ). Serial dilutions of normalized saturated cultures were spotted onto LB agar containing IPTG and incubated at 37°C for approximately 24 h. (b) Ticarcillin-clavulanate plating efficiency assay. Serial dilutions of normalized saturated cultures were spotted onto the surface of Mueller-Hinton agar with (+TIC) or without (-TIC) 4 µg/ml ticarcillin-clavulanate (15:1). Plates were incubated at 37°C for approximately 24 h. (c) A model in which PA3978 inhibits MltF activity, consistent with the Ticarcillin/Clavulanate resistance phenotypes of ΔPA3978 and Δ*mltF* mutants.

A Δ*mltF* mutation suppressed the XcpQ-sensitivity and TIC-resistance phenotypes of a ΔPA3978 mutant (Fig. 3a, b). This suggested that increased MltF activity might be responsible for the ΔPA3978 mutant phenotypes, which would occur if PA3978 was an inhibitor of MltF. Consistent with this hypothesis, Δ*mltF* was epistatic to ΔPA3978, so that a ΔPA3978 mutation had no effect on XcpQ-sensitivity or TIC resistance in a Δ*mltF* strain (Fig. 3a, b). We attempted to test this hypothesis further by determining if increased *mltF* expression phenocopied a ΔPA3978 mutant, but plasmid-encoded *mltF* expression was highly toxic (data not shown). However, in the TIC resistance assay we noticed that in contrast to the approximately 100-fold increased plating efficiency of a ΔPA3978 mutant, a Δ*mltF* mutation alone caused an approximately 10-fold decrease compared to wild type (Fig. 3b). Collectively, these data are consistent with a model in which lack of MltF activity in the Δ*mltF* mutant caused decreased TIC resistance, whereas increased MltF activity in a ΔPA3978 mutant caused increased TIC-resistance (Fig. 3c).

### PA3978 interacts with the catalytic domain of MltF

The structure of *P. aeruginosa* MltF has been solved in both active and inactive conformations (30). It has an N-terminal domain (NTD) with a regulatory role, and a C-terminal domain (CTD) with the catalytic activity (30; Fig. 4a). The switch from inactive to active states occurs when the regulatory NTD is occupied by cell wall-derived muropeptides, which causes a global conformational change that opens the active site in the CTD for catalysis (30). We used the BACTH assay to test if PA3978 interacted with the NTD and/or CTD of MltF. The results indicated that PA3978 interacts with the catalytic CTD of MltF (Fig. 4b).

**FIG 4.**
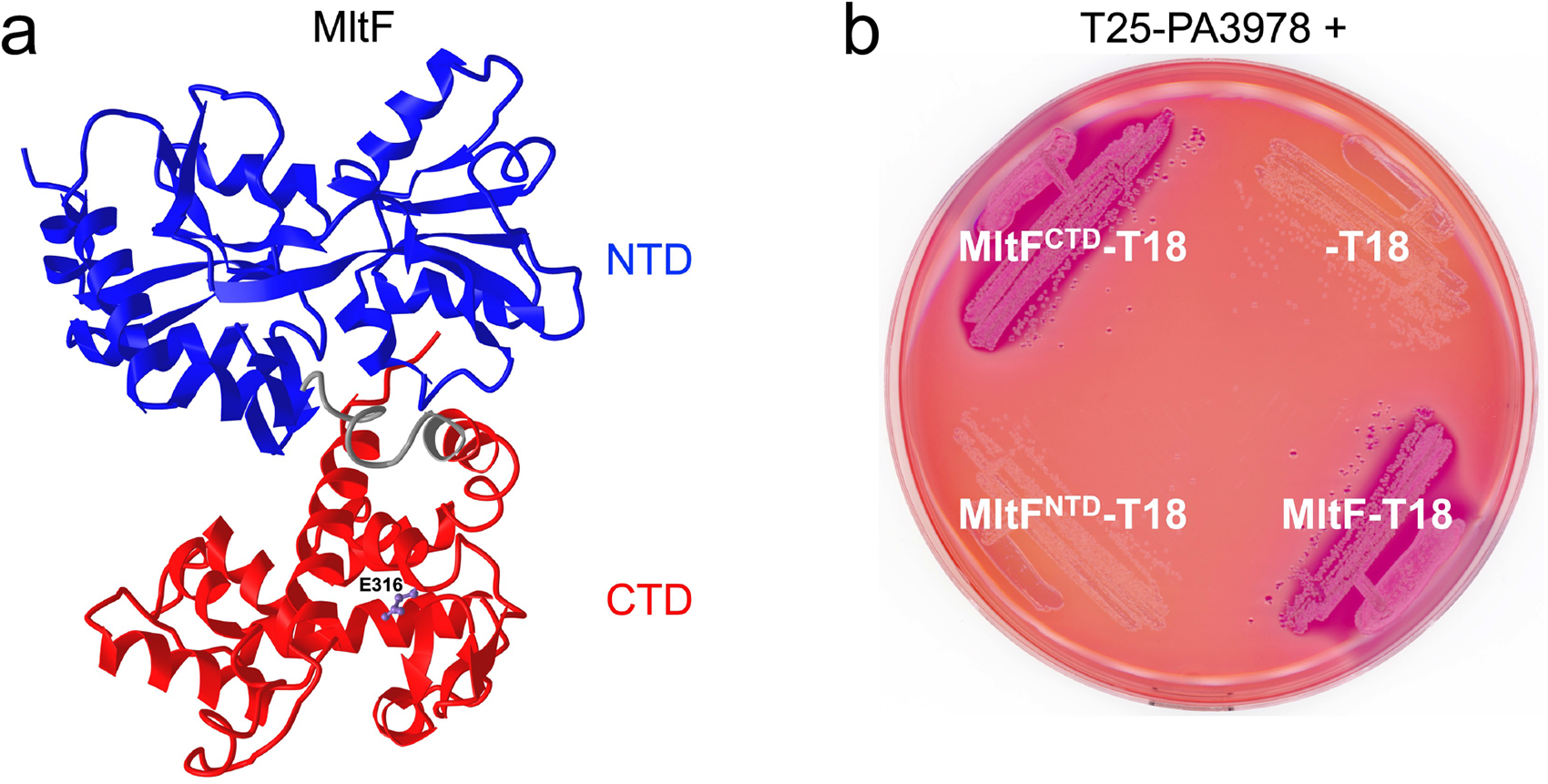
MltF domain analysis. (a) Structure of MltF in its active conformation from the Protein Data Bank (ID: 4P11). The N-terminal regulatory domain is colored in blue, the C-terminal catalytic domain in red, and the linker region between them is gray. The side chain of the catalytic glutamate is in purple and labeled E316. (b) Bacterial two-hybrid analysis. *E. coli* BTH101 strains grown on MacConkey-maltose agar contained a plasmid encoding PA3978 fused to the C-terminus of Cya-T25 and a second plasmid encoding full length MltF, its N-terminal (NTD) or C-terminal (CTD) only, or nothing fused to the N-terminus of Cya-T18 as indicated.

### PA3978 inhibits MltF hydrolase activity

Genetic analysis was consistent with elevated MltF activity in a ΔPA3978 mutant, and the BACTH experiments indicated that PA3978 binds to the catalytic domain of MltF (Figs. 3-4). We hypothesized that PA3978 binds to the catalytic domain of MltF to inhibit its activity. To test this hypothesis, we used a version of the turbidometric assay of Hash, which has been used widely to monitor lysozyme and lytic transglycosylase activities, including for *E. coli* MltF (32–36). This assay monitors the decrease in turbidity of suspended Gram-positive *Micrococcus luteus* cells as their peptidoglycan cell wall is hydrolyzed. The purified C-terminal catalytic domain of MltF (MltF-cat) was active in this assay (Fig. 5a). Activity (reduction in turbidity) was modest, but comparable to that of *E. coli* MltF-cat (36). As a negative control, we tested a version in which the catalytic glutamate was changed to glutamine (E316Q). As expected, the minor decrease in turbidity with this mutant was indistinguishable from that of a *M. luteus* suspension with no enzyme added, which results from cells slowly settling out of suspension over time (Fig. 5a). When PA3978 was added to reactions containing MltF-cat it inhibited the activity, with a molar ratio of 2:1 (PA3978:MltF) reproducibly causing maximum inhibition (Fig. 5a and data not shown). A PA3978-only assay was indistinguishable from *M. luteus* alone, showing that PA3978 has no effect in the absence of MltF-cat (Fig. 5b). PA3978 also inhibited the full length MltF protein, although this version of MltF had very low activity, perhaps because it purified mostly in the inactive conformation, as reported before (Fig. 5b and ref. 30). Finally, although we could not test the effect of PA3978 on MltF2 because MltF2-cat was inactive in the turbidometric assay (data not shown), PA3978 did not inhibit the activity of the lytic transglycosylase RlpA (2:1 ratio PA3978:RlpA; Fig. 5c). PA3978 also had no effect on c-type lysozyme (∼ 40:1 ratio PA3978:lysozyme; Fig. 5c). These *in vitro* data support the hypothesis that PA3978 is a direct inhibitor of MltF.

**FIG 5.**
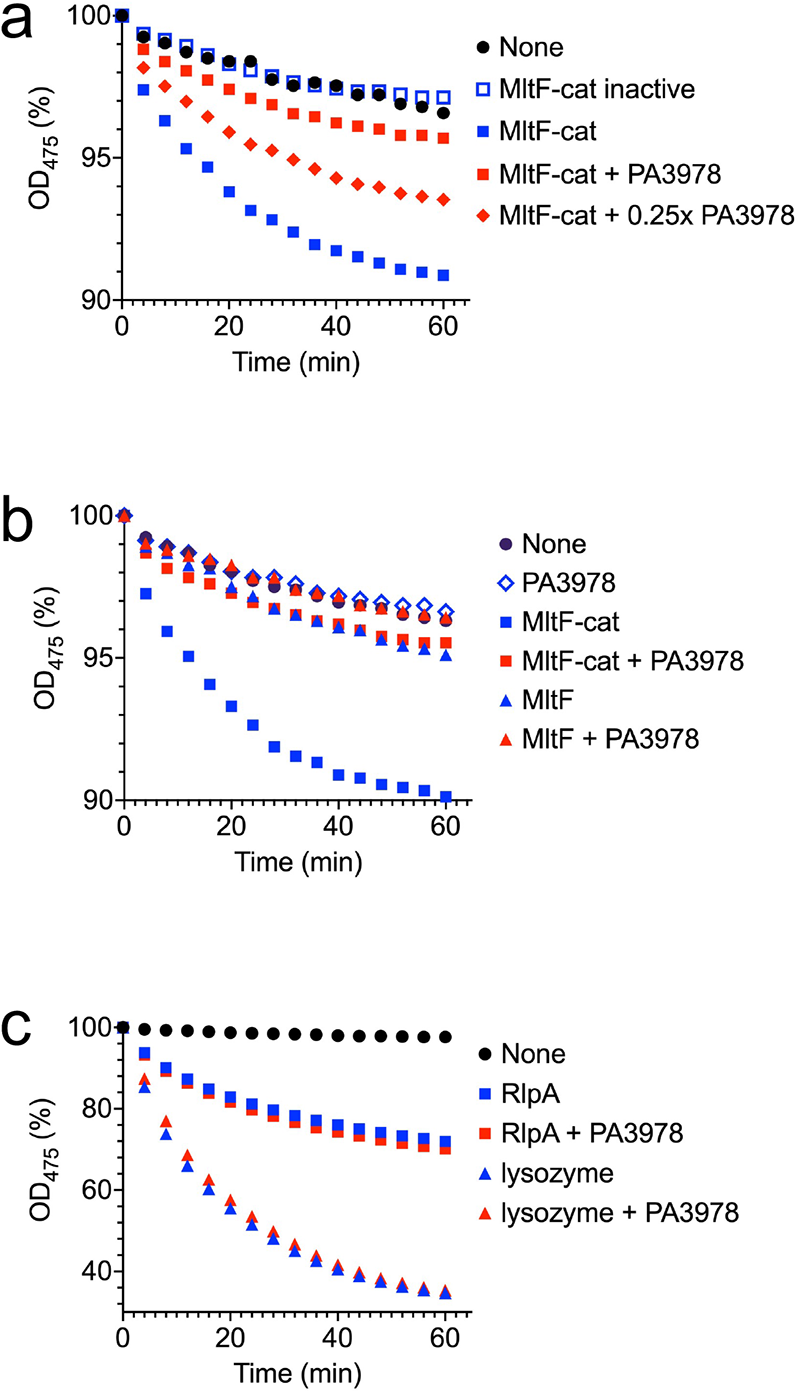
PA3978 inhibits MltF *in vitro*. Turbidometric assay of *M. luteus* hydrolysis. *M. luteus* cells suspended in 0.1 M MOPS, pH 6.5 were incubated with the indicated proteins. The resulting decrease in OD_475_ is shown as a percentage of the initial value for each individual sample. (a) PA3978 inhibits the catalytic domain of MltF. Protein concentrations were 0.44 µM (MltF-cat), 0.88 µM (PA3978) and 0.22 µM (0.25x PA3978). (b) PA3978 inhibits full length MltF. Protein concentrations were 0.44 µM (MltF and MltF-cat) and 0.88 µM (PA3978). (c) PA3978 does not inhibit RlpA or c-type lysozyme. Protein concentrations were 0.44 µM (RlpA), 0.02 µM (c-type lysozyme) and 0.88 µM (PA3978). Data in each individual panel are single representative experiments in which all reactions were run and analyzed simultaneously (conclusions were verified in repeat assays).

### PA5502 is an anti-inhibitor of PA3978

Next, we turned our attention to the other interaction partner of PA3978, the predicted outer membrane lipoprotein PA5502. A ΔPA5502 in frame deletion mutation did not cause a phenotype in the XcpQ-sensitivity assay, but there was an approximately 10-fold decrease in the TIC resistance plating efficiency assay (Fig. 6a). However, a ΔPA5502 mutation had no effect on TIC resistance in ΔPA3978 or Δ*mltF* strains (Fig. 6a). A model consistent with these phenotypes is one in which PA5502 interferes with the inhibitory activity of PA3978 (Fig. 6b). Therefore, in a ΔPA5502 mutant, PA3978 inhibition of MltF increases, reducing MltF activity and decreasing TIC resistance (Fig. 6 a, b). In contrast, with no PA3978 for PA5502 to act on in a ΔPA3978 strain, or no MltF for PA3978 to act on in a Δ*mltF* strain, the removal of PA5502 has no effect on TIC resistance (Fig. 6 a,b).

**FIG 6.**
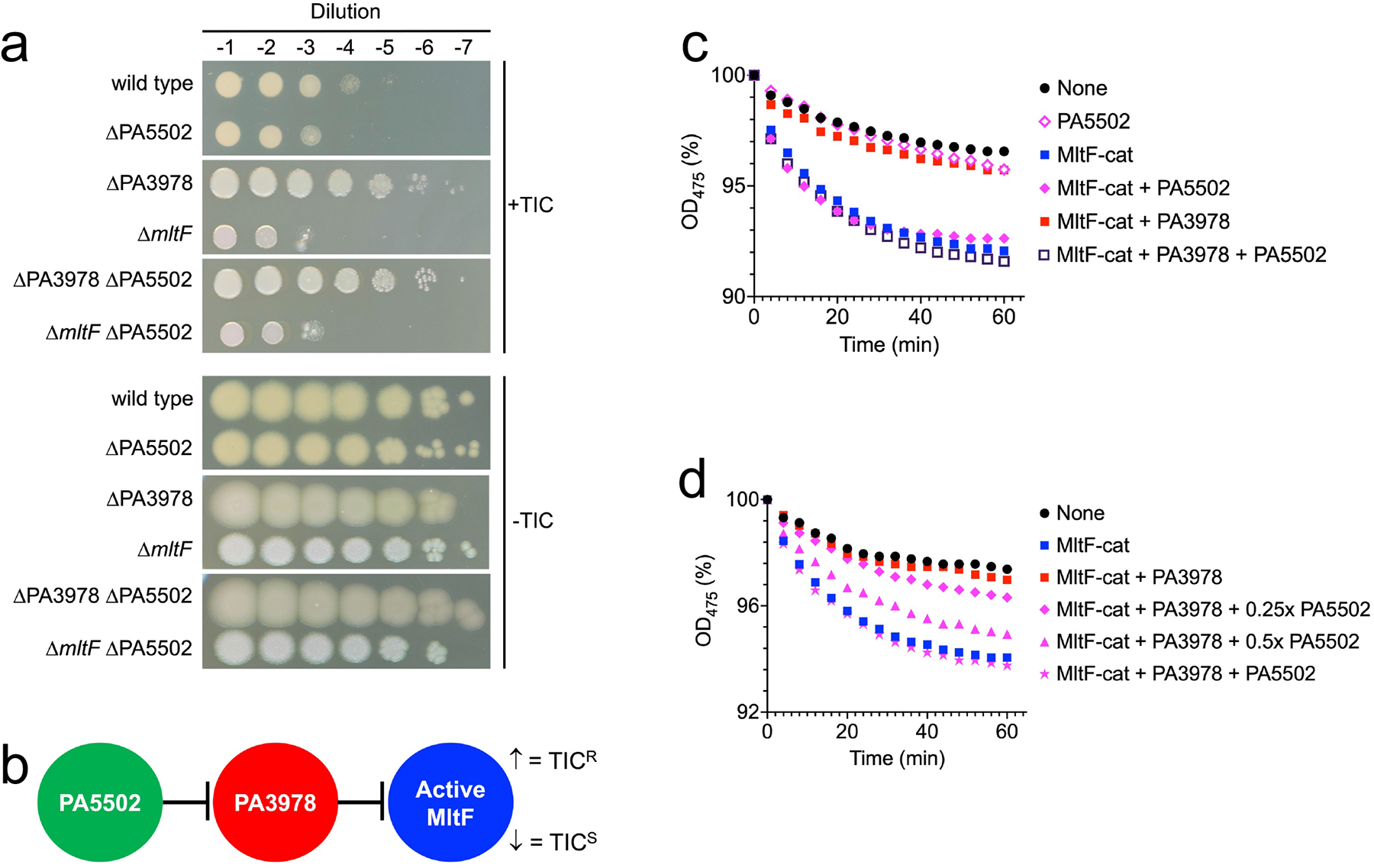
PA5502 is an anti-inhibitor of PA3978. (a) Ticarcillin-clavulanate plating efficiency assay. Serial dilutions of normalized saturated cultures were spotted onto the surface of Mueller-Hinton agar with (+TIC) or without (-TIC) 4 µg/ml ticarcillin-clavulanate (15:1). Plates were incubated at 37°C for approximately 24 h. (b) Model to explain Ticarcillin-clavulanate resistance phenotypes. In a PA3978*^+^ mltF*^+^ strain, ΔPA5502 removes the anti-inhibition activity on PA3978, which increases inhibition of MltF by PA3978 and reduces Ticarcillin-clavulanate resistance. However, in ΔPA3978 or Δ*mltF* strains, ΔPA5502 has no effect on resistance. (c) PA5502 interferes with PA3978-dependent inhibition of MltF *in vitro*. Protein concentrations were 0.44 µM (MltF-cat), 0.88 µM (PA3978 and PA5502). (d) Anti-inhibitory activity of PA5502 is dose dependent. Protein concentrations were 0.44 µM (MltF-cat), 0.88 µM (PA3978 and PA5502), 0.44 µM (0.5x PA5502) and 0.22 µM (0.25x PA5502). For panels (c) and (d) the decrease in OD_475_ of suspended *M. luteus* cells is shown as a percentage of the initial value for each individual suspension. Data in each individual panel are single representative experiments in which all reactions were run and analyzed simultaneously (conclusions were verified in repeat assays).

To test the anti-inhibitor model for PA5502 directly we used the *in vitro* turbidometric assay. Addition of PA5502 only was indistinguishable from an assay with *M. luteus* alone, showing that PA5502 itself is inert in this assay (Fig. 6b). MltF-cat was active, and robustly inhibited by PA3978, providing a third independent confirmation of the earlier reactions in Figure 5a and 5b (Fig. 6c). However, when PA5502 was also added, the inhibition was completely reversed (molar ratios 2:2:1, PA5502:PA3978:MltF). The maximum effect was observed when PA5502 was equimolar to PA3978 (Fig. 6d). Importantly, PA5502 had no affect on MltF-cat activity when it was added to a reaction containing MltF-cat but not PA3978 (Fig. 6c). These data support a model in which PA5502 is an anti-inhibitor of PA3978, rather than a direct activator of MltF (Fig. 6b). This is also consistent with the earlier conclusion that PA5502 binds to PA3978, but not to MltF (Fig. 2d).

### PA3978 interacts with an alpha-helical N-terminal domain of PA5502

AlphaFold has predicted a two-domain organization for PA5502, with an α-helical NTD separated from a globular CTD by an unstructured linker region (Fig. 7a). To determine which of these two PA5502 domains interacts with PA3978, we used the BACTH assay, as we had done earlier for the two domains of MltF (Fig. 4). Similar to the MltF domain analysis, the results for PA5502 were unambiguous, suggesting that PA3978 binds to the α-helical NTD (Fig. 7b). We do not yet know the role of the globular CTD of PA5502, but we speculate that it might be involved in determining when or where PA5502 interacts with PA3978 (see Discussion).

**FIG 7.**
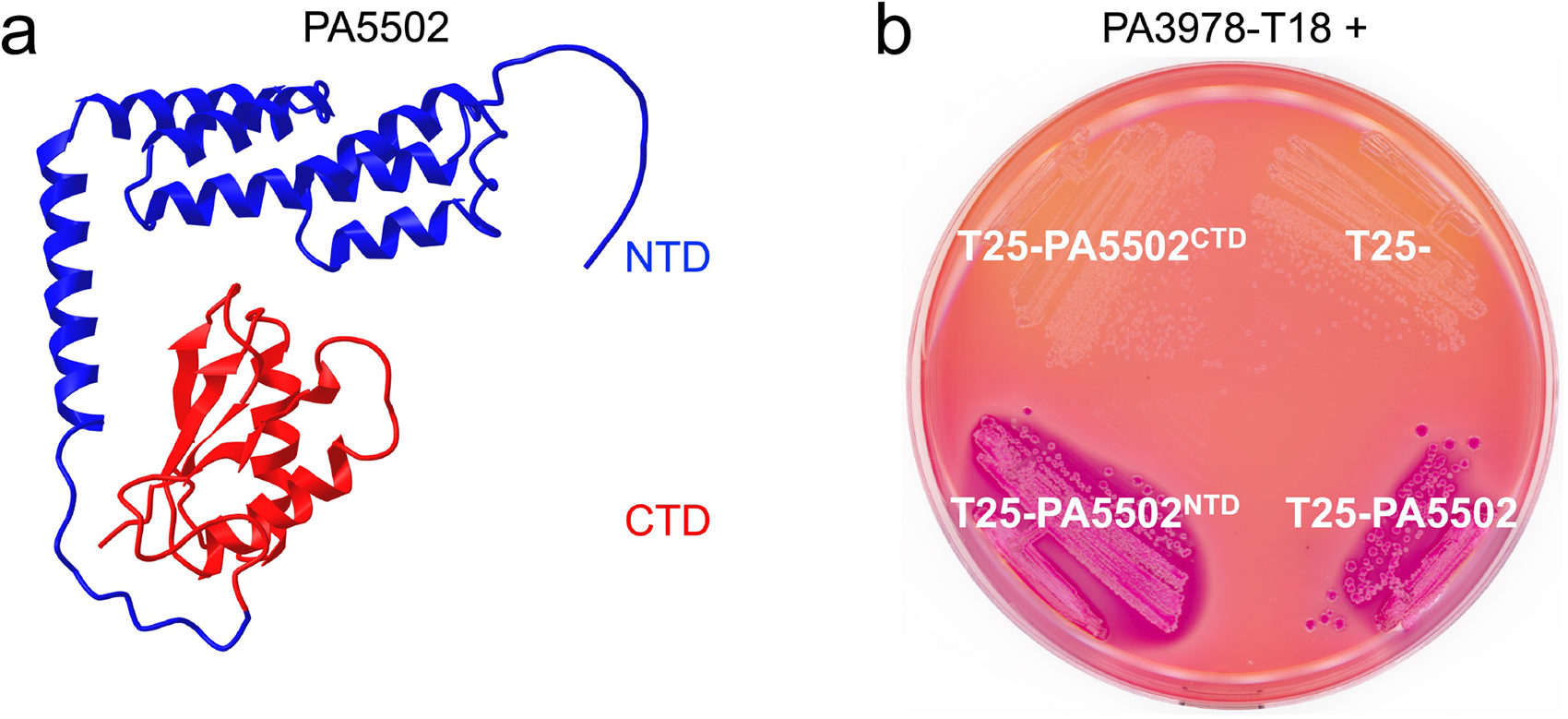
PA5502 domain analysis. (a) AlphaFold-predicted structure of the mature PA5502 protein from the AlphaFold Protein Structure Database. The N-terminal domain is colored in blue and the C-terminal domain in red. (b) Bacterial two-hybrid analysis. *E. coli* BTH101 strains grown on MacConkey-maltose agar contained a plasmid encoding PA3978 fused to the N-terminus of Cya-T18 and a second plasmid encoding full length PA5502, its N-terminal (NTD) or C-terminal (CTD) only, or nothing fused to the C-terminus of Cya-T25 as indicated.

## DISCUSSION

Based on the data presented here, we propose a working model in which two previously uncharacterized proteins function as a regulatory system to control the activity of the lytic transglycosylase MltF (Fig. 8). In addition to bring consistent with the *in vivo* and *in vitro* data from this study, we have also expanded the model to include some speculation that will help to guide future experiments. PA3978 is a direct inhibitor of MltF, and we propose to name this protein Ilt (inhibitor of lytic transglycosylase). PA5502 inhibits this function of Ilt, and for PA5502 we propose the name Lii (lytic transglycosylase inhibitor, inhibitor). Our model implies that it is Lii that might control whether or not Ilt can inhibit MltF. The simplest mechanism for this control to occur is if the binding of Ilt to Lii^NTD^, or to MltF^CTD^, is mutually exclusive (Fig. 8). However, we cannot yet rule out the possibility that Lii interacts with the Ilt-MltF complex and prevents Ilt form inhibiting MltF by some other mechanism. We also speculate that the globular CTD of Lii could be involved in the control mechanism, for example by affecting when or where the Lii^NTD^ binds to Ilt. Using the predicted structure of Lii^CTD^ as the search template with the DALI or PDBeFold servers returned the OmpA-like peptidoglycan-binding domains of some proteins as similar structures (37, 38; data not shown). The Lii^CTD^ primary sequence does not predict this domain, but the same was true for one of the structural homologs, the *Salmonella enterica* peptidoglycan-binding protein SiiA. The similarity of the SiiA CTD to OmpA-like PG-binding domains was discovered from its structure but not from its primary sequence (39). If Lii^CTD^ interacted with peptidoglycan, it could detect a cell wall signal that affects its ability to sequester Ilt, and/or only interact with peptidoglycan in specific locations. The latter possibility might mean that Lii would sequester Ilt only in specific places, allowing MltF to catalyze localized peptidoglycan hydrolysis there. Even if Lii^CTD^ does not interact with peptidoglycan, its role still might be to influence the Lii^NTD^-Ilt interaction by another mechanism. Investigating the function of this domain, along with the precise locations of Lii and Ilt within the cell envelope, is one of the major goals for our future work.

**FIG 8.**
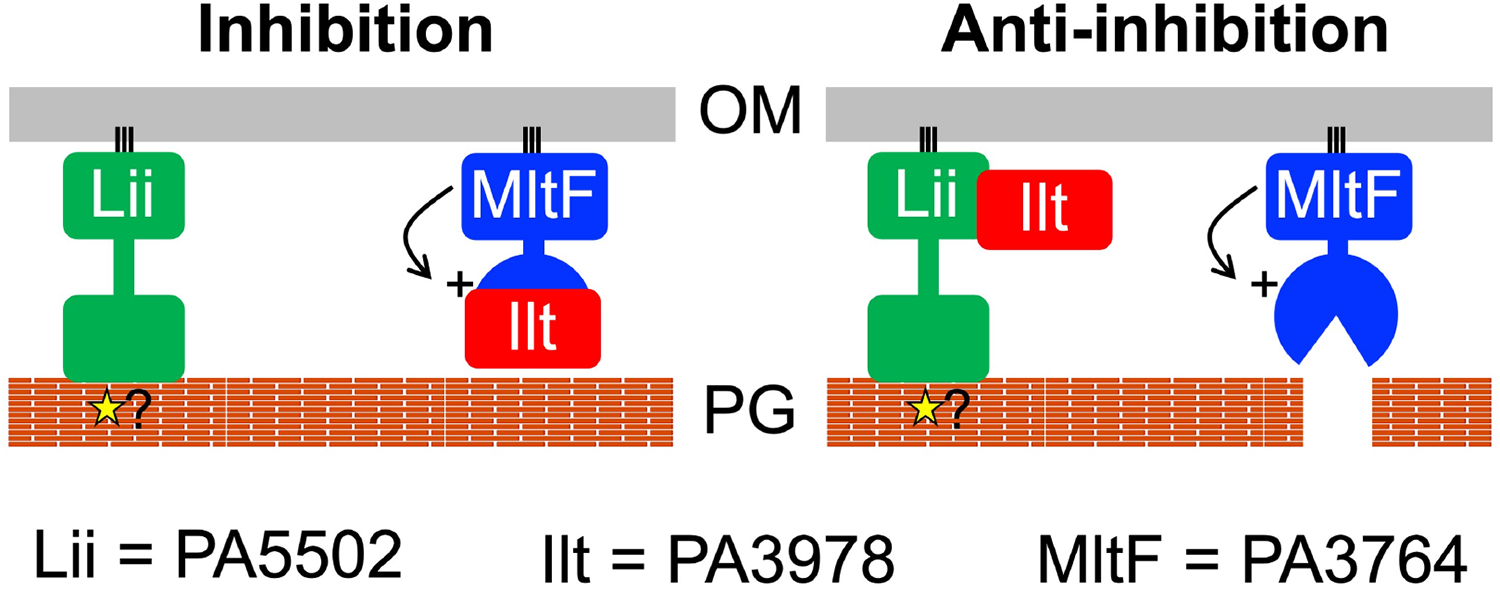
Working model for the Ilt-Lii inhibitor/anti-inhibitor system. Ilt binds to the catalytic domain of MltF and inhibits it from cleaving the cell wall, even if the MltF N-terminal domain-mediated allosteric activation mechanism has occurred (indicated by line arrow and “+” symbol). Lii inhibits this inhibitory function of Ilt by sequestering it away from MltF. The globular C-terminal domain of Lii might play a role in controlling when the Lii^NTD^-Ilt interaction occurs, perhaps by interacting with peptidoglycan. For example, it could detect a cell wall signal (star symbol) that affects its ability to bind to Ilt, and/or only interact with peptidoglycan in specific places. The latter possibility would mean that Lii sequesters Ilt at only those places, allowing MltF to catalyze localized peptidoglycan hydrolysis. Alternatively, the Lii^CTD^ might influence when the Lii^NTD^-Ilt interaction occurs by a mechanism that does not involve peptidoglycan binding, or have a different role altogether. OM = outer membrane; PG = peptidoglycan.

The role of PA3978 as a lytic transglycosylase inhibitor is similar to a group of bacterial proteins known as lysozyme inhibitors, which are also found in *P. aeruginosa* (32). The first was discovered in *E. coli*, named Ivy (inhibitor of <u>v</u>ertebrate lysozyme), and was shown to inhibit c-type lysozyme (40). Later, another group of lysozyme inhibitors was discovered, called the MliC/PliC family (<u>m</u>embrane bound/<u>p</u>eriplasmic lysozyme inhibitors of c-type lysozyme) (32). Since then, bacterial inhibitors of other lysozyme types were also found (41, 42). *P. aeruginosa* has two Ivy proteins, IvyP1/PA3902 and IvyP2/PA5481, as well as MliC/PA0867 (33). As their name indicates, originally these proteins were proposed to protect bacteria from host lysozyme during infection. There is some evidence to support this role in Gram negative bacteria, despite them having an outer membrane barrier to lysozyme (e.g., 32, 41-43). However, later it was shown that IvyP1 and IvyP2 of *P. aeruginosa* can inhibit both c-type lysozyme and MltB (the only lytic transglycosylase tested), leading to the proposal that the true function of these proteins might be to control the activity of bacterial lytic transglycosylases. In other words, they might play a similar role to Ilt. However, there are some key differences between Ilt and the lysozyme inhibitors. Ilt did not inhibit c-type lysozyme (Fig. 5c), its predicted structure is not homologous to any of the bacterial lysozyme inhibitors, and it is acted on by an anti-inhibitor protein (Lii), which has not been described for any of the lysozyme inhibitors. Therefore, Ilt and Lii provide a novel example of a direct control mechanism of lytic transglycosylase activity.

Initially, we were surprised to discover a new control mechanism for MltF, because one had already been described. Structural analysis revealed that *P. aeruginosa* MltF occurs in active and inactive conformations, with a switch to the active conformation being triggered when the regulatory NTD of MltF is engaged by cell wall-derived muropeptides (30). The fact that MltF has this allosteric control mechanism in response to the level of a cell wall component raises the question of what the purpose of the Lii-Ilt system might be. We propose that Ilt acts as an overriding off switch that prevents MltF from degrading peptidoglycan, even if the allosteric mechanism has activated it. Our data support this hypothesis. First, a Δ*ilt* mutant has MltF-dependent phenotypes consistent with increased MltF activity, suggesting that its function is to inhibit catalytically active MltF *in vivo* (Fig. 3). Second, Ilt interacts with the catalytic domain of MltF, which might allow it to occlude the substrate-binding region and active site (Fig. 4b). Third, the activity of full length MltF was inhibited by Ilt *in vitro*, which further indicates that Ilt can inhibit MltF in its active conformation (Fig. 5b). Therefore, we speculate that the role of the Lii-Ilt system might be to modulate MltF in specific conditions and/or at specific locations within the cell envelope. In other words, it might facilitate a localized peptidoglycan hydrolysis role for MltF. It could also be significant that Ilt and MltF co-purified with similar abundance according to mass spectrometry, whereas Lii co-purified with much less abundance (Fig. 2b). When coupled with the occurrence of robust MltF-dependent phenotypes in a Δ*ilt* mutant, this could indicate that much of the MltF in the cell is being inhibited by Ilt. This model would fit with the proposal that Lii might sequester only a fraction of the cellular Ilt away from MltF, and do so at specific times and/or locations.

We found an *ilt* null mutant in a screen for sensitivity to overproduction of the XcpQ secretin, which was the same screen in which we discovered a *ctpA* null mutant (15, 16). Four of the five known CtpA proteolytic substrates are predicted peptidoglycan cross-link hydrolases (14). Therefore, like Ilt, CtpA is an inhibitor of cell wall hydrolase activity, albeit a different family of hydrolases, and with inhibition in this case occurring by hydrolase degradation. We do not yet know the molecular mechanism that explains the secretin-sensitivity phenotype of the *ilt* and *ctpA* null mutants. However, one thing that both mutants have in common is likely to be increased peptidoglycan hydrolysis. As mentioned in the Introduction, this might increase secretin-toxicity by disturbing secretin trafficking, or by impacting phospholipid and lipopolysaccharide biosynthesis, which share precursors in common with peptidoglycan. Regardless of the mechanism, this raises the possibility that altered peptidoglycan, and perhaps peptidoglycan hydrolysis specifically, might also be linked to some other mutations that we had found in the original screen, including several genes of unknown function (16; unpublished data). Exploring this possibility is one of our goals for the future.

The *mltF* (PA3764), *ilt* (PA3978) and *lii* (PA5502) genes are not linked, which raises the possibility that Lii might have functions beyond controlling Ilt, and/or Ilt might have functions beyond inhibiting MltF. We currently have no data to support those possibilities, and several observations argue against it. First, the XcpQ-sensitivity and TIC resistance phenotypes of an *ilt* null mutant were completely suppressed by Δ*mltF* (Fig. 3). We have also noticed that an *ilt* null mutant often has a slightly different color and more surface spreading on agar, which was also suppressed by Δ*mltF* (e.g., Fig. 3b). Second, in our original Ilt-FLAG co-purification assay, Lii and MltF were the only proteins that were both abundant and highly enriched (Fig. 2b). Third, BACTH analysis suggested that Ilt does not interact with at least two other lytic transglycosylases, including another MltF family member that has been named MltF2 (Fig. 2d). Fourth, Ilt did not affect the activities of c-type lysozyme or the lytic transglycosylase RlpA *in vitro* (Fig. 5c; we could not reach a conclusion for inhibition of MltF2 *in vitro* because it was inactive in the turbidometric assay). Despite these negative data, the possibility of other functions for Ilt and Lii must remain open until comprehensive analyses are done.

A homolog of Ilt in *Acinetobacter baylyi*, which was named Gcf, was found in a screen for mutations that blocked giant cell formation resulting from the inhibition of peptidoglycan biosynthesis (44; GenBank accession CAG67453.1). The mature Gcf and Ilt proteins are similar in size, both have three Sel-1 like/TPR repeats, they are 37% identical, and 65% similar (data not shown). An *mltD* null mutant was found in the same screen as *gcf*, the *mltD* and *gcf* null mutants had indistinguishable phenotypes, and the mutations were epistatic to each other. This led the authors to speculate that Gcf might activate MltD. The *gcf* and *mltD* phenotypes were also recapitulated in *Acinetobacter baumannii* (44). Although the speculation that Gcf might activate MltD was not tested experimentally, these observations raise the possibility that Ilt homologs might modulate lytic transglycosylase activity in many bacteria. It will be interesting to investigate if Ilt can increase the activity of any lytic transglycosylases in *P. aeruginosa*, in addition to its role as an inhibitor of MltF.

In summary, we have assigned roles to two previously uncharacterized cell envelope proteins in *P. aeruginosa*. PA3978, which we have named Ilt, interacts with and inhibits MltF. PA5502, which we have named Lii, interacts with and inhibits this role of Ilt. Some of our future goals include investigating if these roles of Ilt and Lii are regulated, determining the function of the CTD of Lii and whether or not it influences the Lii-Ilt interaction, and exploring the possibility that Ilt and Lii might have other roles/targets, including any other lytic transglycosylases, of which there are eleven in *P. aeruginosa* (31). We will also be interested to determine if any of the other genes of unknown function that we found in our secretin-sensitivity screen also affect the cell wall, and the activity of cell wall hydrolases in particular. Identifying new peptidoglycan hydrolase control mechanisms, such as the one we have reported here, also has the potential to provide new targets for therapeutic intervention.

## MATERIALS AND METHODS

### Bacterial strains and standard growth conditions

*P. aeruginosa* strains and plasmids used in this study are listed in Table 1. *E. coli* K-12 strain SM10 was used for conjugation of plasmids into *P. aeruginosa* (45). For routine propagation, bacteria were grown in Luria-Bertani (LB) broth, composed of 1% (w/v) tryptone, 0.5% (w/v) yeast extract, 1% (w/v) NaCl, or on LB agar. To select for *P. aeruginosa* exconjugants after mating with *E. coli* donor strains, bacteria were recovered on Vogel-Bonner minimal agar with appropriate antibiotics (46).

**TABLE 1.**
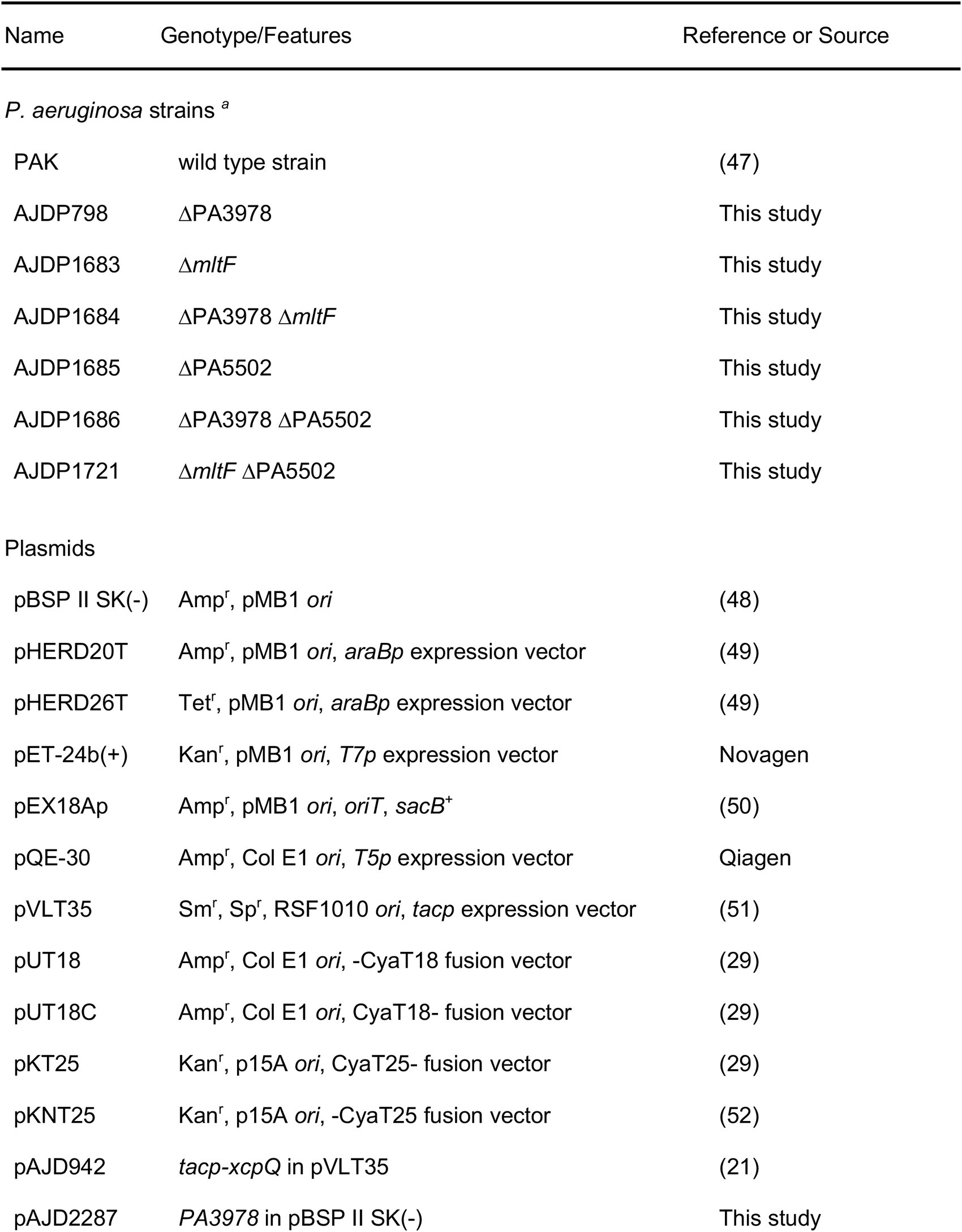
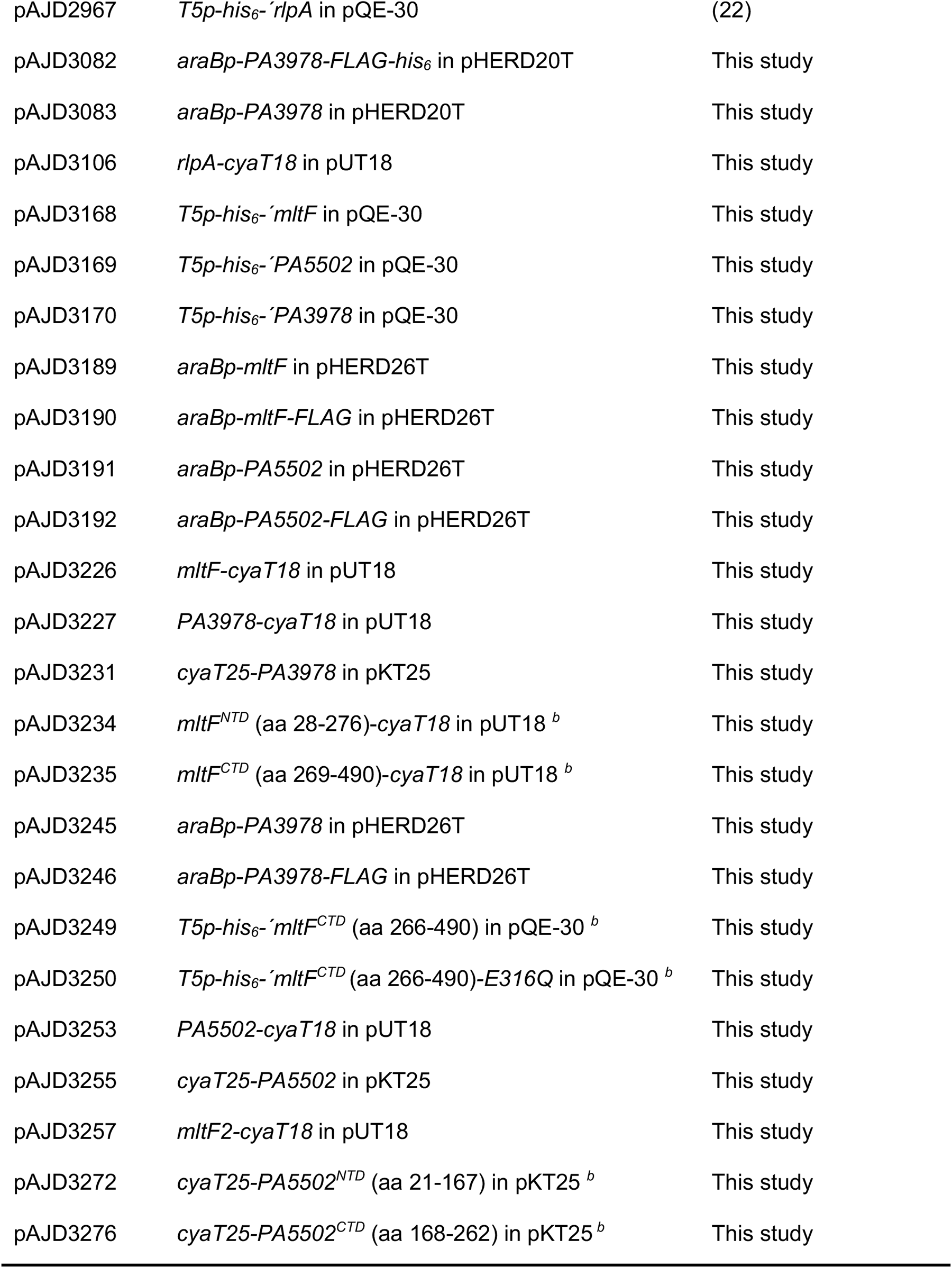
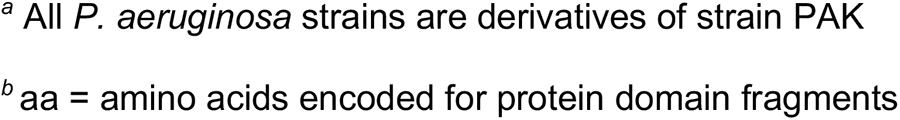
Strains and plasmids.

### Plasmid and strain constructions

To construct strains with ΔPA3978, Δ*mltF* or ΔPA5502 in frame deletion mutations, fragments of ∼0.55 kb each corresponding to regions upstream and downstream of the deletion site were amplified by PCR and cloned into pEX18Ap. These plasmids were integrated into the *P. aeruginosa* chromosome after conjugation from *E. coli*, and sucrose-resistant, carbenicillin-sensitive segregants were isolated on LB agar containing 10% sucrose (44). Deletions were verified by PCR analysis of genomic DNA.

Plasmids encoding C-terminal epitope-tagged proteins were constructed by amplifying the genes from *P. aeruginosa* PAK DNA using a downstream primer that included a region encoding the tag, followed by a stop codon. For plasmids encoding proteins without any added tags, the downstream primer annealed immediately downstream of the stop codon. The amplified fragments were cloned into pHERD20T or pHERD26T using restriction sites incorporated by the PCR primers. Plasmids encoding proteins with N-terminal His_6_ tags were constructed by amplifying the region encoding the proteins or protein domain (but without any N-terminal signal sequence) from *P. aeruginosa* PAK DNA, using a downstream primer that annealed immediately after the stop codon. These fragments were cloned into pQE-30 using restriction sites incorporated by the PCR primers.

For BACTH assay plasmids, gene fragments without regions encoding signal sequences were amplified by PCR and cloned into the two-hybrid vectors as XbaI-KpnI fragments, except for *rlpA*, which was a XbaI-BglII fragment (restriction sites were introduced by the PCR primers).

### XcpQ sensitivity assay

Strains were grown to saturation at 37°C in LB broth containing 250 µg/ml streptomycin and 500 µg/ml spectinomycin. Optical densities at 600 nm (OD_600_) were determined and then cultures were diluted to OD_600_=1.0 in LB broth. 3 µl of undiluted and serial 10-fold dilutions (10^-1^ – 10^-6^) of each sample were spotted onto the surface of LB agar (30 ml in a standard 100 mm petri dish) containing 250 µg/ml streptomycin, 500 µg/ml spectinomycin, and 1 mM IPTG. Plates were incubated at 37°C for approximately 24 h.

### Ticarcillin-clavulanate sensitivity assays

Strains were grown to saturation at 37°C in LB broth, OD_600_ was determined and then cultures were diluted to OD_600_=1.0 in LB broth. For disc diffusion assays, a cotton swab was used to spread the diluted culture onto Mueller-Hinton agar (60 ml in a large 150 mm diameter petri dish). Becton Dickinson Sensi-Discs with Ticarcillin-clavulanic acid (75/10 µg) were then laid on the surface and the plates were incubated at 37°C for approximately 24 h. For spot dilution assays, 3 µl of serial 10-fold dilutions (10^-1^ – 10^-7^) of each sample were spotted onto the surface of Mueller-Hinton agar (30 ml in a standard 100 mm petri dish) containing 4 µg/ml ticarcillin-clavulanate (15:1, Alfa Aesar). Plates were incubated at 37°C for approximately 24 h.

### Anti-FLAG co-immunoprecipitation assay

Bacteria were inoculated into 100 ml of LB broth in a 250 ml flask to an OD_600_ of 0.05, and grown at 37°C with aeration for 5 h. Equivalent amounts of bacterial cells from all strains were collected by centrifugation, washed in 10 mM Tris-HCl, 10% (w/v) glycerol, pH 7.5 and resuspended in 2 ml lysis buffer (50 mM Tris-HCl, 300 mM NaCl, 10% (w/v) glycerol, pH 7.5). Roche complete protease inhibitors were added, cells were frozen, disrupted by sonication, and then 1% (w/v) Lauryldimethylamine N-oxide (LDAO) was added, followed by incubation with rotation for 1 h at 4°C. Insoluble material was removed by centrifugation at 16,000 x*g* for 30 min at 4°C. 35 µl of anti-FLAG M2 affinity resin (SigmaAldrich) in lysis buffer was added to the supernatant and incubated for 2 h at 4°C with rotation. A 1-ml spin column (Pierce; 69725) was used to wash the resin five times with 500 µl lysis buffer containing 0.1% (w/v) Triton X-100 and 10% (w/v) glycerol, and then five times with 500 µl lysis buffer containing 0.1% (w/v) Triton X-100 only. Proteins were eluted by addition of 100 µl of 400 µg/ml 3x FLAG peptide in lysis buffer containing 0.1% (w/v) Triton X-10, and incubation with rotation at 4°C for 30 min. Proteins present in these samples were identified by liquid chromatography-mass spectrometry (NYU School of Medicine Proteomics Laboratory) or analyzed by immunoblotting.

### PA3978 polyclonal antiserum production and immunoblotting

*E. coli* strain M15 [pREP4] (Qiagen) containing plasmid pAJD3179 encoding His_6_-PA3978 was grown in LB broth to mid-log phase at 37°C with aeration. Protein production was induced with 1 mM IPTG for 3 h at 37°C. The protein was purified under denaturing conditions by nickel-nitrilotriacetic acid (NTA)-agarose affinity chromatography as described by the manufacturer (Qiagen). A polyclonal rabbit antiserum was raised by Labcorp Early Development Laboratories Inc. Specificity was verified by detection of a protein of the predicted size in a whole cell lysate of a *P. aeruginosa* ΔPA3978 mutant containing a PA3978 expression plasmid, but not in a strain with the empty vector control, as well as by analysis of whole cell lysates of *E. coli* DH5α transformants with the same plasmid pair.

Samples separated by SDS-PAGE were transferred to nitrocellulose by semi dry electroblotting. Chemiluminescent detection followed incubation with the PA3978 polyclonal antiserum described above, or anti-FLAG M2 monoclonal antibody (Sigma), then goat anti-rabbit IgG (Sigma) or goat anti-mouse IgG (Sigma) horseradish peroxidase conjugates used at the manufacturers recommended dilution.

### Bacterial two-hybrid assay

Pairs of plasmids encoding Cya-T25 and Cya-T18 derivatives were introduced simultaneously into *E. coli* Δ*cya* mutant strain BTH101 (Euromedex) by calcium chloride transformation. Transformants were streaked onto MacConkey-maltose agar and incubated at 30°C for approximately 40 h.

### Protein purification for lytic activity assays

*E. coli* strain M15 [pREP4] (Qiagen) containing a plasmid pQE-30 protein-encoding derivative was grown in LB broth to mid-log phase at 37°C with aeration. Protein production was induced with 1 mM IPTG for 3 h at 37°C. Most proteins were insoluble when overproduced in *E. coli* and were purified using a denaturation-renaturation procedure. Cells were resuspended by stirring for 1 h in 6 M GuHCl, 0.1 M NaH_2_PO4, 0.01 M Tris-Cl, 5 mM Imidazole, 1% (w/v) Triton X-100, pH 8.0. The suspension was sonicated and insoluble materials were removed by centrifugation at 10,000 x*g* for 30 min. NTA-agarose was added to the supernatant followed by stirring for 1 h. The suspension was poured into a drip column to collect the NTA-agarose, then washed with 15 NTA-agarose volumes of 100 mM NaH_2_PO4, 10 mM Tris-HCl, 1% (w/v) Triton X-100, 20 mM imidazole, 8M Urea, pH 8.0. It was then washed with 25 column NTA-agarose volumes of 50 mM NaH_2_PO4 pH 8.0, 300mM NaCl, 20mM Imidazole for protein renaturation. Proteins were eluted in 2 ml fractions of 50 mM NaH_2_PO4 pH 8.0, 300 mM NaCl containing increasing imidazole concentrations (50 – 250 mM, in 50 mM increments). One exception to this procedure was full length MltF, which was purified under entirely native conditions because it was completely inactive if first denatured. Cells were resuspended in 50mM NaH_2_PO4, 300 mM NaCl, 10 mM imidazole, pH 8.0 and incubated with 1 mg/ml lysozyme for 30 min on ice. Cells were then disrupted by sonication and insoluble materials were removed by centrifugation. NTA-agarose was added to the supernatant followed by stirring for 1 h. The NTA-agarose was collected with a drip column, then washed with 40 NTA-agarose volumes of 50mM NaH_2_PO4, 300 mM NaCl, 10 mM imidazole, pH 8.0, followed by 40 NTA-agarose volumes of the same buffer but with 20 mM imidazole. Proteins were eluted as described above.

### *In vitro* lytic enzyme activity assay

This was a version of the turbidometric assay of Hash, which has been used to monitor lysozyme and lytic transglycosylase activities (32–36). Protein samples were incubated with *M. luteus* cells (0.4 mg/ml of purchased *Micrococcus lysodeikticus* cells; Sigma-Aldrich M3770) in a total volume of 1 ml of 0.1 M MOPS, pH 6.5. A reaction containing bovine serum albumin as the only protein served as a negative control to monitor decreased turbidity resulting from the settling of the suspended *M. luteus* cells. For each individual figure panel, all protein sample additions were of the same total volume and in the same elution buffer (either 150 mM or 200 mM Imidazole). Also, all reactions within each individual figure panel were run and analyzed simultaneously in a multi-sample carousel of a Beckman-Coulter DU-720 spectrophotometer. Optical density of each sample at 475 nm was monitored every 4 min for 1 hour.

### Data availability

All data for the proteomic mass spectrometry analysis are available at the Mass Spectrometry Interactive Virtual environment (MassIVE), a part of ProteomeXchange. DOI: Pending.

## ACKNOWLEDGEMENTS

Research was supported by the National Institute of Allergy and Infectious Diseases (NIAID) of the National Institutes of Health, under Award Number R01AI136901. The content is solely the responsibility of the authors and does not necessarily represent the official views of the National Medicine’s proteomics laboratory was partly supported by NYU Grossman School of Medicine. We thank Maureen Ty for constructing strain AJDP798 and plasmid pAJD2287, Dolon Chakraborty for constructing plasmid pAJD3106, and Alexis Sommerfield for experimental advice. We also thank Heran Darwin, Kévin Rome and Alexis Sommerfield for comments on a draft version of the manuscript.

---

**Supplementary Figure S1.**
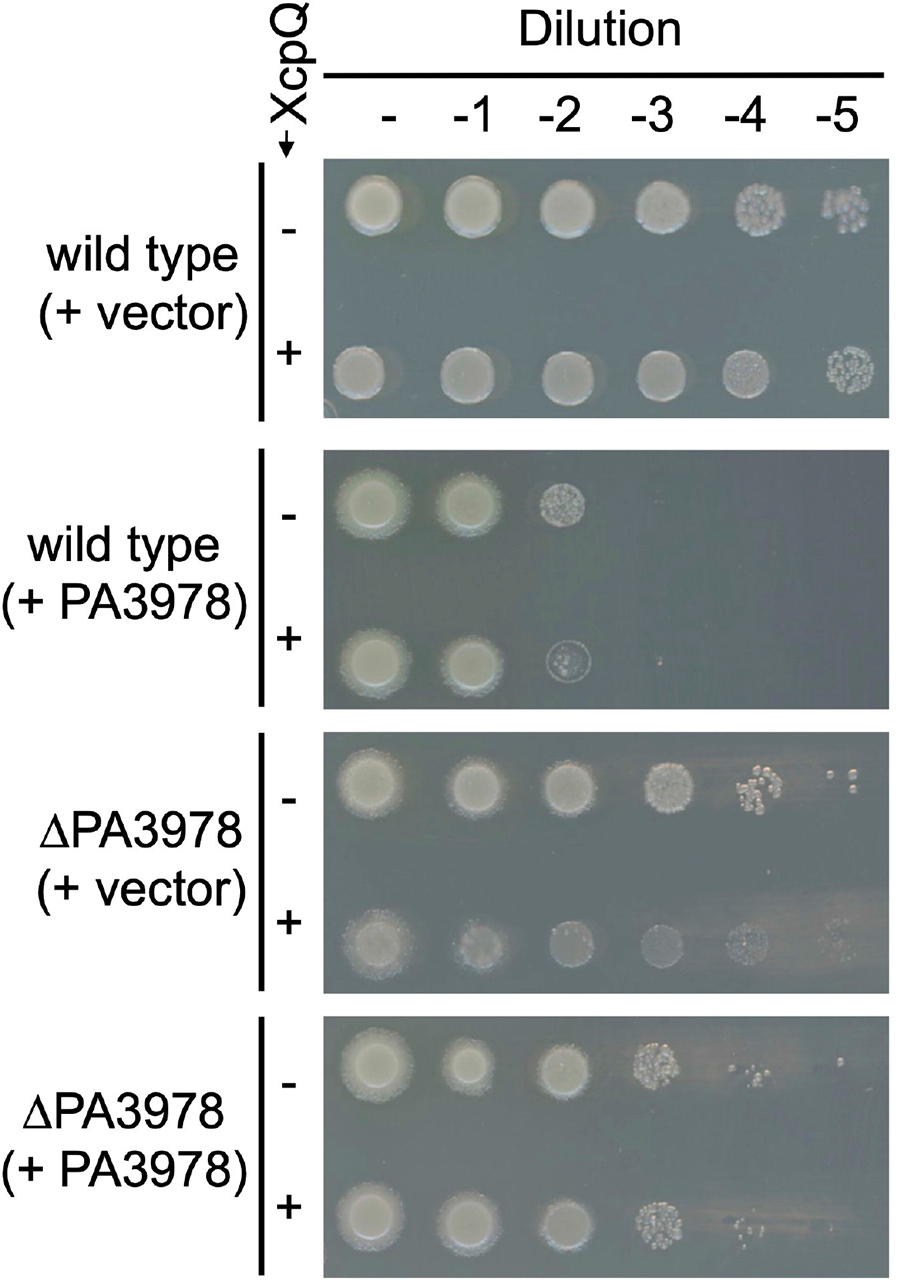
Complementation of ΔPA3978 mutant XcpQ-sensitivity. Wild type and ΔPA3978 strains contained plasmid pAJD2287 encoding PA3978 expressed from its native promoter, or the empty vector control. Strains also contained either the *tac* promoter expression plasmid pVLT35 (-XcpQ), or the *xcpQ*^+^ derivative pAJD942 (+XcpQ). Serial dilutions of normalized saturated cultures were spotted onto LB agar containing 125 µM IPTG and incubated at 37°C for approximately 24 h. Plasmid pAJD2287 was toxic, especially in the wild type strain that also had endogenous PA3978. Therefore, XcpQ production was induced with a lower IPTG concentration in this experiment to reduce the overall amount of stress.

**Supplemental Table S1:**
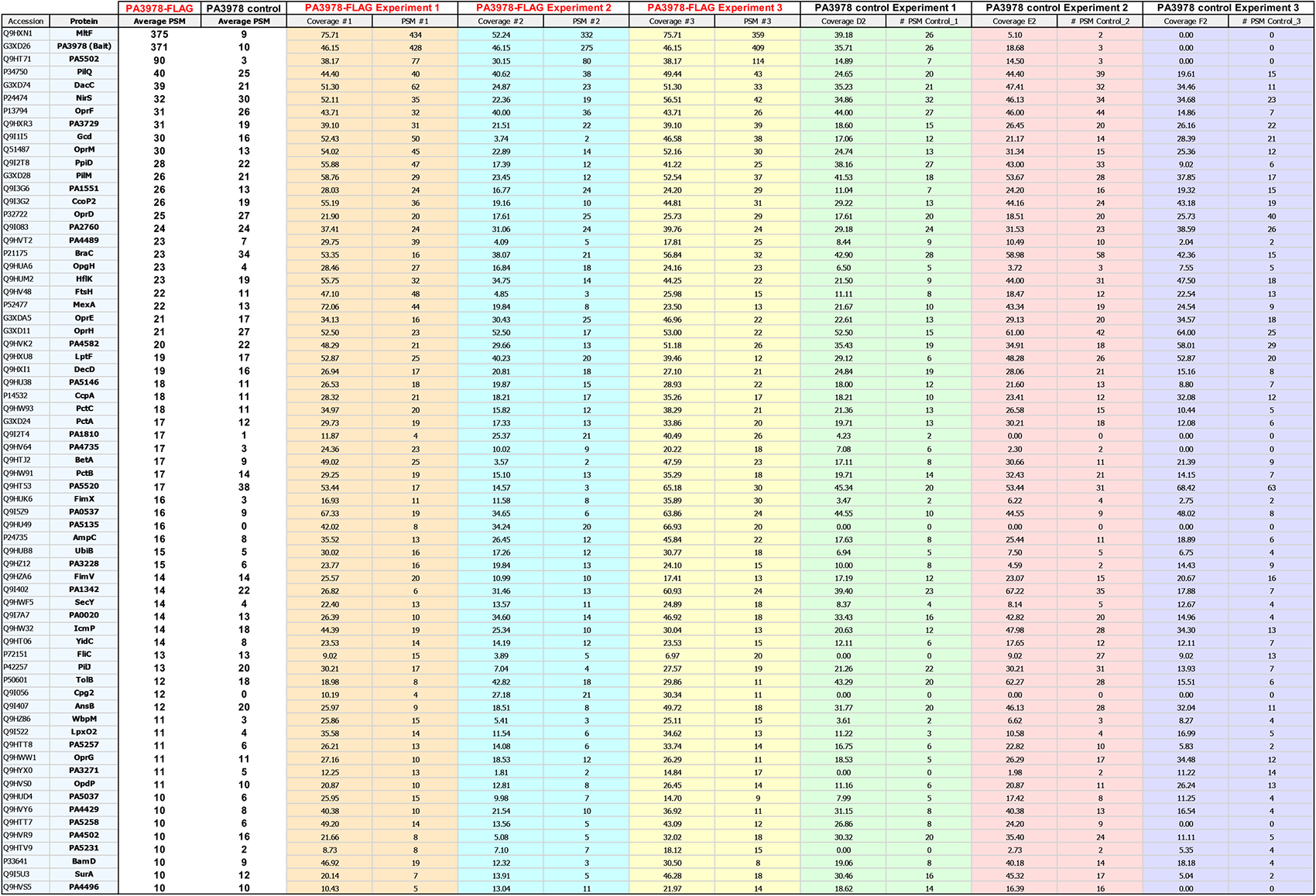
Known or predicted cell envelope proteins that co-purified in each of three PA3978-FLAG pulldowns with an average PSM ≥ 10.

